# Evolutionary diversification reveals distinct somatic versus germline cytoskeletal functions of the Arp2 branched actin nucleator protein

**DOI:** 10.1101/2023.02.25.530036

**Authors:** Kaitlin A. Stromberg, Tristan Spain, Sarah A. Tomlin, Kristen Dominique Amarillo, Courtney M. Schroeder

**Affiliations:** Department of Pharmacology, UT Southwestern Medical Center, Dallas, TX; Division of Basic Sciences, Fred Hutchinson Cancer Research Center, Seattle, WA; Howard Hughes Medical Institute, Fred Hutchinson Cancer Research Center, Seattle, WA

**Keywords:** cytoskeleton, actin, Arp2, evolution, fertility, sperm development

## Abstract

Branched actin networks are critical in many cellular processes, including cell motility and division. Arp2, a protein within the 7-membered Arp2/3 complex, is responsible for generating branched actin. Given its essential roles, Arp2 evolves under stringent sequence conservation throughout eukaryotic evolution. We unexpectedly discovered recurrent evolutionary diversification of Arp2 in *Drosophila*, yielding independently arising paralogs *Arp2D* in *obscura* species and *Arp2D2* in *montium* species. Both paralogs are unusually testis-enriched in expression relative to Arp2. We investigated whether their sequence divergence from canonical Arp2 led to functional specialization by replacing *Arp2* in *D. melanogaster* with either *Arp2D* or *Arp2D2*. Despite their divergence, we surprisingly found both complement Arp2’s essential function in the soma, suggesting they have preserved the ability to polymerize branched actin even in a non-native species. However, we found that *Arp2D*-expressing males are subfertile and display many defects throughout sperm development. We pinpointed two highly diverged structural regions in Arp2D that contribute to these defects: subdomain 2 and the C-terminus. We expected that germline function would be rescued by replacing Arp2D’s long and charged C-terminus with Arp2’s short C-terminus, yet surprisingly, the essential somatic function of Arp2D was lost. Therefore, while Arp2D’s structural divergence is incompatible with *D. melanogaster* sperm development, its unique C-terminus has evolved a critical role in actin polymerization. Our findings suggest canonical Arp2’s function differs between somatic versus germline contexts, and Arp2 paralogs have recurrently evolved and specialized for actin branching in the testis.

## Introduction

The Arp2/3 complex is a highly conserved protein complex that nucleates branched actin networks and is found in almost all eukaryotes. The complex comprises seven proteins, consisting of two actin-related proteins, Arp2 and Arp3, and five other subunits, ArpC1-5. The complex docks onto a mother actin filament and promotes the nucleation of a daughter filament at a 70° angle from the mother filament^1^. Arp2/3-generated branched actin networks are often found at the surface of cell membranes and are vital for many cellular processes, such as cell motility and endocytosis^2-4^. These actin networks are mechanosensitive and respond to environmental cues by undergoing actin reorganization, implicating Arp2/3 in events such as cellular signaling and energetics^5^. This complex also plays roles in genomic stability^6^ and cell division, particularly in proper chromosome segregation, spindle organization, and mitotic progression^6^. Finally, Arp2/3 also resides in the nucleus, where it aids in DNA damage repair^7,8^.

In addition to its many somatic roles, Arp2/3 has specialized roles in the germline. In *Drosophila*, Arp2/3 is required for generating actin structures found in ovaries^9^ and testes^10^. *Drosophila* ovaries have actin structures known as ring canals, which are critical for successful oogenesis and allow cytoplasmic flow to the oocyte^9^. The Arp2/3 complex localizes to ring canals and is necessary for their expansion in late oogenesis^9^. *Drosophila* sperm development also depends on Arp2/3^11,12^. *Drosophila* sperm develop together within cysts sharing a cytoplasm; mature sperm must be separated following their elongation and tail development^10^. To accomplish this individualization process, an actin cone forms around each sperm within the cyst, and all cones move synchronously along the tail from the basal to the apical end of the testis^10,11^. The Arp2/3 complex localizes to the front of the cones, and branched actin polymerization aids in motility^12^.

Recent studies have begun to reveal the structure and regulation of the Arp2/3 complex in mechanistic detail. Like actin, Arp2 and 3 contain four subdomains with an ATP-binding fold^13^. Upon docking onto the mother actin filament, Arp2/3 undergoes a conformational change from inactive to active, in which Arp2 and Arp3 are no longer splayed but are aligned together, like actin monomers in a filament^14-16^. This re-arrangement of Arps 2 and 3 allow them to become the first two subunits of the daughter actin filament^15,17^. Arp2/3 cannot nucleate actin filaments on their own^18^; instead, nucleation-promoting factors (NPFs) activate Arp2/3 downstream of many signaling pathways^16^. Numerous regulatory factors terminate polymerization and disassemble filaments, replenishing the monomeric actin pool for future nucleation events^19^. Filament nucleation, polymerization, termination, and disassembly are regulated by a myriad of proteins to fine-tune the timing and structural stability of branched actin networks^19^, thus contributing to the diversity of Arp2/3’s functional roles.

Eukaryotic genomes show genetic and functional diversification of the Arp2/3 complex. Humans encode two isoforms of Arp3 (Arp3 and Arp3B), ArpC1 (ArpC1A and ArpC1B), and ArpC5 (ArpC5 and ArpC5L)^20,21^. Arp2/3 complexes containing ArpC1B and ArpC5L differ in their rates of actin polymerization from those containing ArpC1A and ArpC5^22^. Although Arp3B functions similarly to canonical Arp3 in actin polymerization, it diverges from Arp3 in the rate of actin network disassembly^21^. Like humans, *Drosophila* also exhibits diversification of the Arp2/3 complex. *D. melanogaster* encodes two isoforms of Arpc3 (Arpc3a and Arpc3b). Arpc3a is expressed in all tissues, whereas Arpc3b appears to be expressed only in the ovary and localizes to intercellular bridges known as actin rings^9^. Furthermore, an Arp2 paralog, known as Arp2D, emerged in one clade of *Drosophila* species. Arp2D is highly expressed in the male germline and localizes to branched actin networks in the front of actin cones^23^. Arp2/3’s key regulators also show diversification, allowing for differential regulation of Arp2/3 between cell types. Defects in the Arp2/3 regulatory system are implicated in disease^19,22^. Therefore, despite Arp2/3’s stringent conservation, multiple members of the Arp2/3 complex and its regulators have undergone recurrent gene duplication and diversification across phyla, suggesting that evolution is selecting for different ‘flavors’ of Arp2/3 for specialized roles.

Here, we have discovered that recurrent diversification of Arp2 in *Drosophila* has led to functional specialization in the male germline. In addition to *Arp2D* found in the *Drosophila obscura* clade^*23*^, we found a second, independently arising gene duplication of *Arp2* in the *montium* clade, named *Arp2D2*. Both Arp2D and Arp2D2 are exclusively expressed in the male germline of their respective species. Despite their striking sequence divergence, we found that Arp2D and Arp2D2 can polymerize actin in somatic tissues in the non-native species *D. melanogaster* and complement canonical Arp2’s essential function in the soma. However, the more divergent Arp2D paralog cannot complement Arp2 function in sperm development, resulting in many testis defects. Arp2D diverges most from canonical Arp2 at the C-terminus and a loop found in subdomain 2. We found that both regions contribute to defects in germline function, yet the unique C-terminus of Arp2D is essential for its somatic function. These data reveal that evolution has shaped the Arp2 structure for specialized roles in spermatogenesis.

## Results

### Recurrent invention of testis-expressed Arp2 paralogs in *Drosophila* species

The seven-member Arp2/3 complex is present and evolves under stringent sequence constraints in most eukaryotes. However, the subunit Arp2 has undergone unexpected diversification via gene duplication. We previously discovered that an *Arp2* paralog, *Arp2D*, arose via retroduplication in the *obscura* clade and has been retained for over 14 million years^23^. By examining additional sequenced *Drosophila* genomes, we found that *D. kikkawai*^*24*^ and *D. serrata*^*25*^, species of the *montium* clade, also encoded an *Arp2* gene duplication found on chromosome 2 (Figure 1A). We named the newly discovered duplicate ‘*Arp2D2*.’ We queried the *Arp2D2* locus by PCR in 18 additional *montium* species, whose genomes were unsequenced. Together with recently published *montium* genomes^26^, we concluded that *Arp2D2* is present in a shared syntenic location in 27 *montium* species with no exceptions (Figure 1B, Figure S1A, Data S1-3). Based on this, we conclude that *Arp2D2* arose in the common ancestor of the *montium* group at least 15 million years ago^27^ (Figure S1A). Our phylogenetic analysis based on a nucleotide alignment shows that *Arp2D2* sequences form a monophyletic clade that is a well-supported sister clade to the *montium Arp2* orthologs (100% bootstrap support), further confirming *Arp2D2* duplicated from canonical *Arp2* in the common ancestor of the *montium* group (Figure 1B). *Arp2D* orthologs form a separate clade with its closest outgroup being *obscura* Arp2 sequences, supporting independent origins of the *Arp2* paralogs (Figure 1B). *Arp2* encodes six introns (Figure 1A), yet 26 of the 27 *Arp2D2* sequences have none (the *D. diplacantha Arp2D2* subsequently acquired one intron (Figure S1B, Data S2)). Therefore, our phylogenomic analyses reveal that *Arp2D2* and *Arp2D* both arose via independent retroduplication events in which Arp2’s spliced mRNA was reverse transcribed and inserted into the genome^*23*^ (Figure 1A-B). Subsequent to their origin, both *Arp2D* and *Arp2D2* have been retained throughout the *obscura* and *montium* clades, respectively, suggesting that they must confer a functional advantage to their native species.

**Figure 1:**
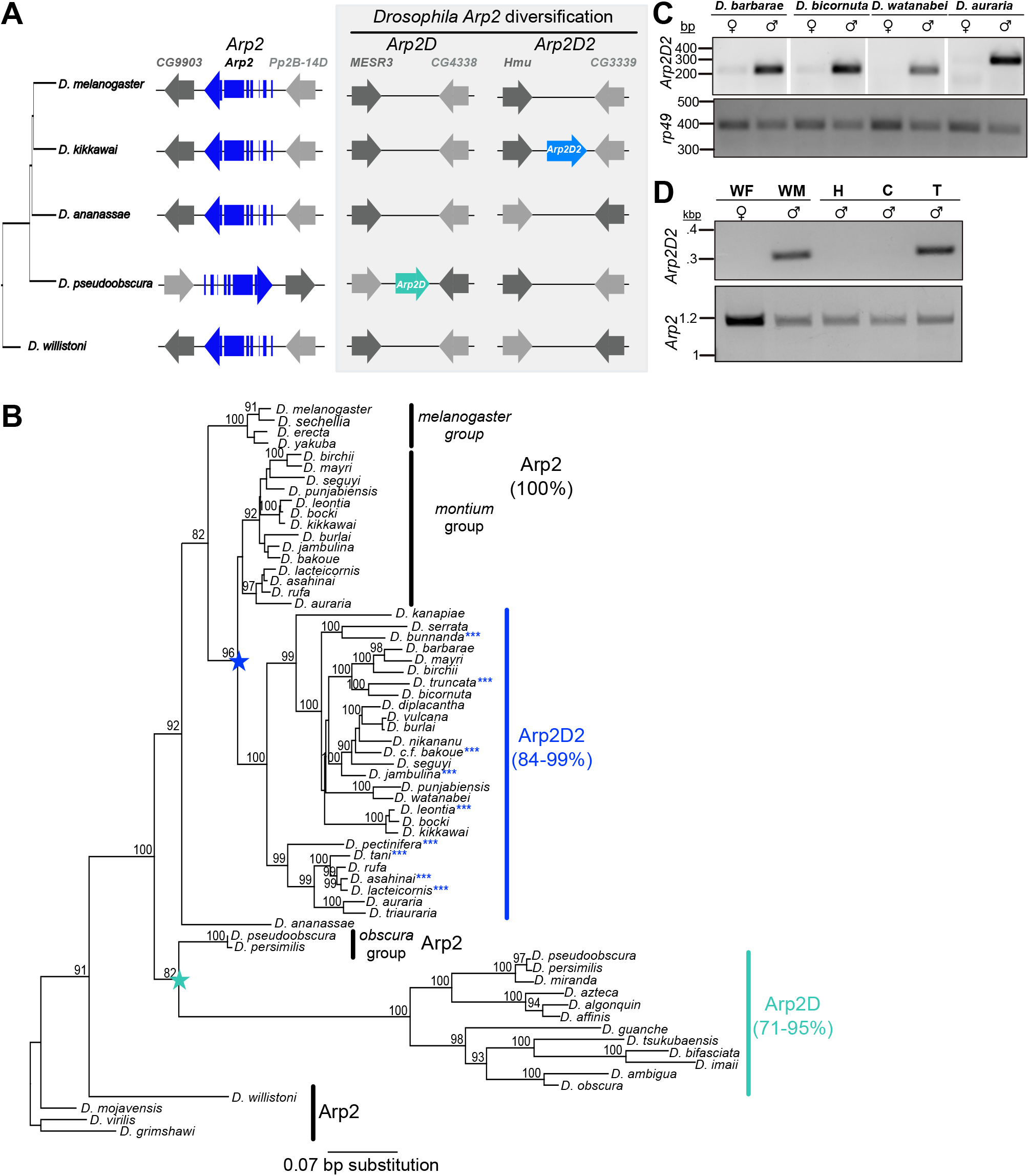
Arp2D2 is a male germline-enriched Arp2 paralog found in the *montium* group species of *Drosophila*. **A)** The syntenic loci of canonical *Arp2* in representative *Drosophila* species spanning 50 million years of evolution. *Arp2D* is found in the *obscura* clade and was described previously^23^. *Arp2D2* was discovered in *montium* group species. **B)** Phylogenetic tree derived from *Arp2, Arp2D*, and *Arp2D2* coding sequence with 100X resampling. Species with blue asterisks have genomes that have been fully sequenced^26^. The ranges of protein sequence identity among the orthologs are given, and nodes with >80% bootstrap support are shown. Stars indicate independent evolutionary origins; *Arp2D2* in the *montium* clade shares a common ancestor with *montium Arp2*; *Arp2D* in the *obscura* clade shares a common ancestor with *obscura Arp2*. **C)** RT-PCRs comparing mRNA expression of *Arp2D2* between females and males in four *montium* species. *Rp49* was used to confirm similar quantities of cDNA among the samples. **D)** RT-PCRs comparing mRNA expression of *Arp2D2* in *D. auraria* whole females (WF), whole males (WM), heads (H), carcasses (C) or testes (T). *Arp2* was used as a control for presence of cDNA among the samples.

Although independently derived from canonical *Arp2, Arp2D* and *Arp2D2* may have converged on similar sequence features associated with their testis-specific roles. However, we observed no conserved residue changes common to all Arp2D and Arp2D2 orthologs but distinct from canonical Arp2, suggesting the two duplicates had divergent evolutionary paths (Figure S1C). Moreover, when we viewed the conserved residue changes on the surface of the Arp2 structure (where protein interactions occur), we did not find residue changes concentrating in one structural region, indicating no ’hot spot’ of diversification for Arp2D or Arp2D2 (Figure S1C).

*Arp2D* exhibits testis-enriched expression, unlike the ubiquitously expressed canonical Arp2^23^. We investigated whether *Arp2D2* also exhibits sex-biased expression. We compared the expression of *Arp2D2* mRNA between males and females in the *montium* species *D. barbarae, D. bicornuta, D. watanabei*, and *D. auraria* by RT-PCR analyses. We detected *Arp2D2* only in males (Figure 1C, Figure S1D-E, Data S1). To determine if expression is localized to the testis, we extracted RNA from dissected *D. auraria* testes, separating them from the head and remaining male carcass. We detected *Arp2D2* mRNA solely in the testis (Figure 1D, Figure S1F-G, Data S1). Thus, two independent gene duplications of *Arp2* exhibit testis-enriched expression, suggesting recurrent *Arp2* diversification may have specifically acquired roles in sperm development.

### Divergent Arp2 duplicates localize to actin in *D. melanogaster*

Our previous study revealed that Arp2D, under the control of its endogenous promoter, localizes to actin in the testis of its native species, *D. pseudoobscura*^23^. However, it was unclear whether it could also do so in heterologous species like *D. melanogaster*. To investigate this possibility, we studied whether divergent *D. pseudoobscura* Arp2D (from the *obscura* group) or *D. auraria* Arp2D2 (from the *montium* group) could localize to actin in *D. melanogaster*. We generated *D. melanogaster* fly lines that express the paralogs tagged with superfolder GFP (sfGFP) at their C-termini (Figure 2A), where canonical Arp2 can be tagged without a noticeable impact on function^28^. We expected the paralogs to be expressed in both germ cells and somatic cells because they are under the control of the endogenous Arp2 promoter (Figure 2A). Indeed, we found that Arp2D, Arp2D2, and canonical Arp2 from *D. pseudoobscura* were highly expressed and localized to actin cones in the testis, even in the presence of endogenous *D. melanogaster* Arp2 (Figure 2C). Actin cones are unique testis-specific actin structures that are specialized for *Drosophila* sperm development^10^. In flies, germ cells remain interconnected throughout development until they are fully mature, and actin cones facilitate the separation of mature sperm^10^ (Figure 2B). Like canonical Arp2, which localizes to the front half of the cone to help produce the cone’s typical fan shape, we observed Arp2D and Arp2D2 also localizing to the cone’s front half (Figure 2C). Based on their localization, we conclude that Arp2D and Arp2D2 fold and localize to actin, suggesting they incorporate into the *D. melanogaster* Arp2/3 complex despite their sequence divergence.

**Figure 2:**
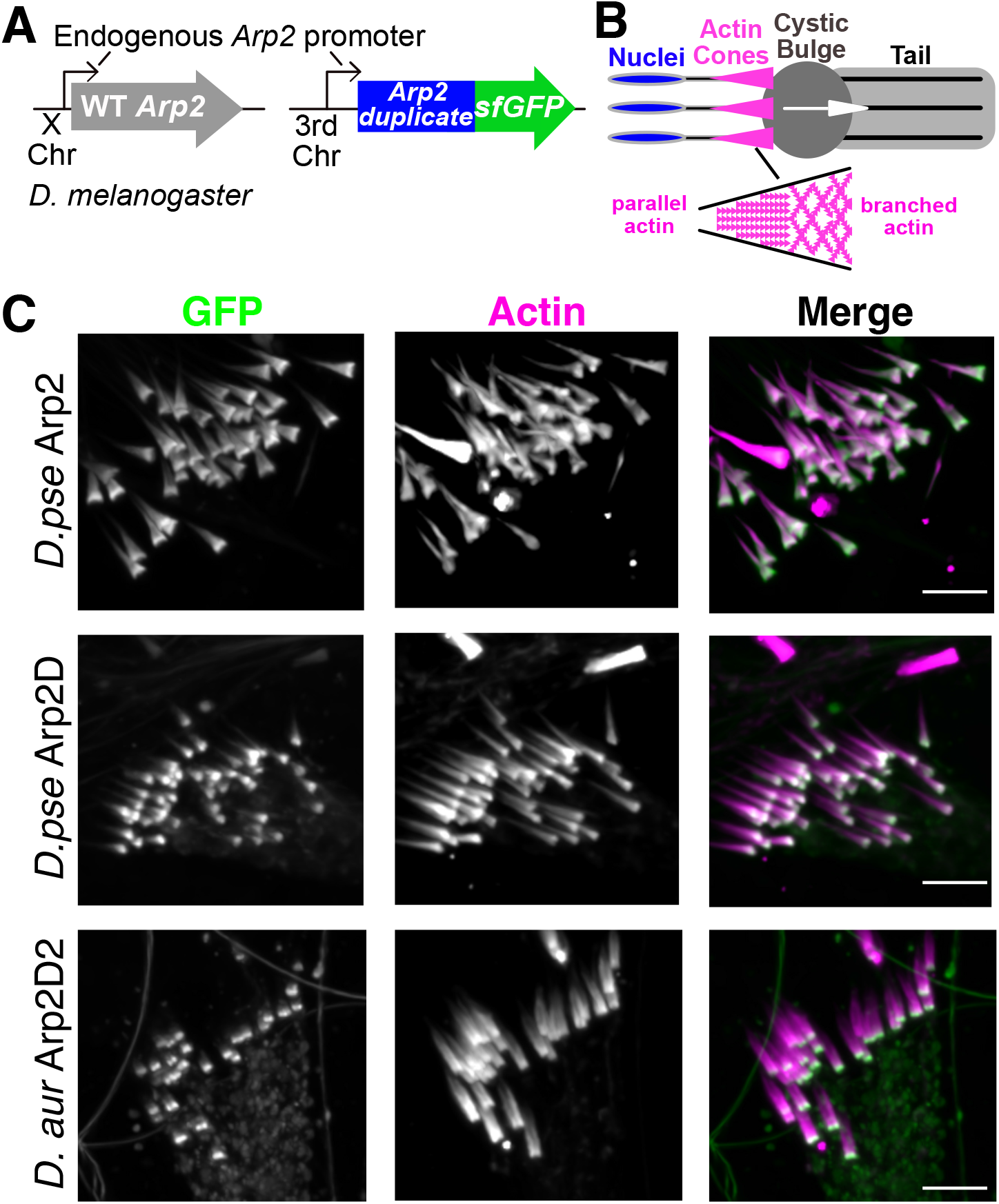
Divergent Arp2 paralogs localize to actin in *D. melanogaster*. **A)** A schematic showing how wildtype *D. melanogaster* flies were genetically modified. Canonical Arp2 (*D. pseudoobscura*), Arp2D (*D. pseudoobscura*) or Arp2D2 (*D. auraria*) were C-terminally tagged with superfolder GFP and inserted on the third chromosome, whereas canonical *Arp2* is on the X-chromosome. All transgenes are under the control of the endogenous *Arp2* promoter. **B)** Schematic depicting individualization of mature sperm with actin cones. Each sperm has one cone, and all cones move synchronously from the sperm head to the end of the sperm tail, moving excess cytoplasm in the ‘cystic bulge.’ Cones contain parallel actin filaments in the rear and Arp2/3-generated branched actin networks in the front half. **C)** Live imaging of individualizing sperm from lines in (A). Manual separation of cysts for improved staining and imaging led to scattered cones. Native GFP fluorescence of the tagged proteins was imaged with actin. The merge of actin (magenta) and Arp2 (green) is white. Scale bars are 10 μm.

### Divergent Arp2 duplicates rescue actin polymerization *in vivo*

Mutations or knockdowns of Arp2/3 complex components lead to lethality^3,9^. We next tested if the *Arp2D* and Arp2D2 paralogs could rescue the loss of *D. melanogaster Arp2*. For this, we first generated an *Arp2-KO* fly line by deleting the entire gene and replacing it with an eye-expressed *DsRed* cassette using CRISPR/Cas9 technology (Figure 3A). Upstream to *DsRed* in the KO locus, we introduced an *attP* site, which we subsequently used for targeted transgenesis (Figure 3A). As expected, our *Arp2*-KOs exhibited homozygous lethality. Because Arp2 is on the X chromosome, only heterozygous *Arp2-KO* females survived, whereas hemizygous *Arp2-KO* males did not.

**Figure 3:**
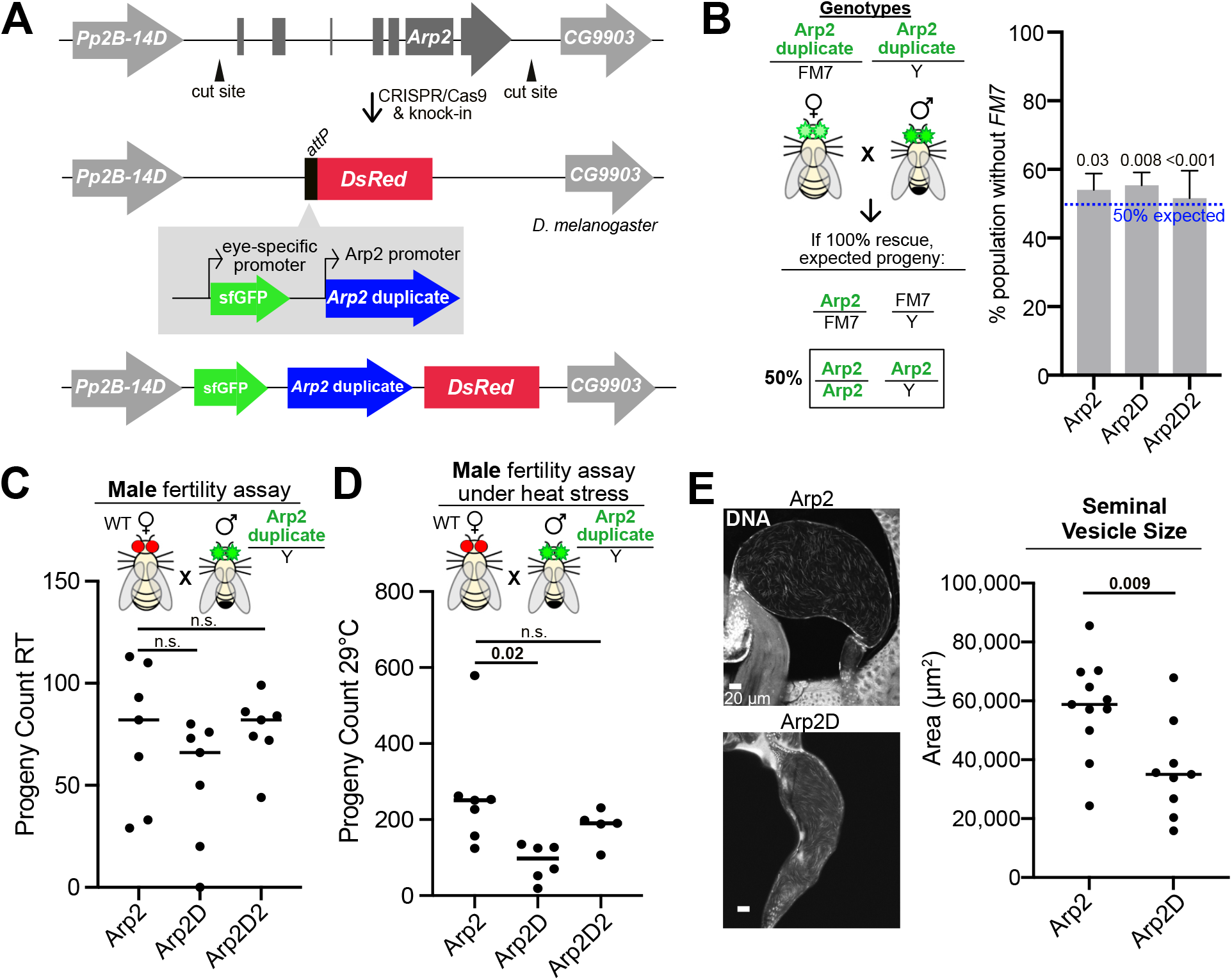
Arp2 paralogs rescue Arp2 somatic functions, yet Arp2D-expressing males are subfertile. **A)** Schematic of CRISPR strategy used to generate Arp2 knockouts (KOs). Cut sites were in the intergenic regions upstream and downstream of canonical *Arp2. DsRed*, under the control of an eye-specific promoter, was inserted to track the KO allele. An *attP* site is upstream of *DsRed* and was leveraged for site-directed transgenesis. Canonical *D. melanogaster Arp2, D. pseudoobscura Arp2D*, or *D. auraria Arp2D2* were inserted at the *attP* site, and superfolder GFP was used as an eye marker to track the transgene. All transgenes were intronless and tagless. **B)** Left, schematic of crossing scheme used to test for rescue of *Arp2*-KO lethality. Heterozygous females (transgene balanced with *FM7*) were crossed to hemizygous males. Eye color indicates presence of the transgene, with heterozygous females having dimmer GFP-positive eyes. Right, percent of the progeny population without *FM7* (homozygous females and hemizygous males) is shown for canonical *Arp2, Arp2D*, and *Arp2D2*. Each cross had 6-8 replicates. Genotype fractions were compared to Mendelian expectation using a chi-squared test. Homozygotes were higher than expected, suggesting a slight fitness cost from the balancer. **C)** A male fertility assay was conducted by crossing males (generated in A) to wildtype females (Oregon R). Arp2D-expressing males exhibited a reduction, though not significant) at room-temperature. **D)** A male fertility assay as done in (C) except under heat stress (29°C). Arp2D-expressing males produced significantly less progeny than canonical Arp2- or Arp2D2-expressing males. **E)** Left, seminal vesicles from Arp2-and Arp2D-expressing males from (A) imaged with a DNA probe (20 μm scale bar). Right, area was measured and compared, revealing significantly smaller seminal vesicles from Arp2D-expressing males (non-virgin).

We next inserted *D. pseudoobscura Arp2D, D. auraria Arp2D2*, or canonical *D. melanogaster* Arp2 (as a positive control) in the *Arp2*-KO locus to test for rescue. We used intronless *D. melanogaster* Arp2 because *Arp2D* and *Arp2D2* lack introns, and we wanted to prevent splicing differences from leading to any phenotypes. All transgenes were introduced using the same *attP* site; they all possess the same upstream and downstream ∼1kb regions of *D. melanogaster* canonical Arp2, which included the 5’- and 3’ UTRs (Figure 3A). Because Arp2D is only 70% identical to canonical Arp2, we expected Arp2D would fail to rescue lethality, whereas Arp2D2, being 96% identical, was likely to rescue. We were surprised to find that we were able to obtain homozygous females and hemizygous males expressing Arp2D or Arp2D2, suggesting both rescued the *Arp2*-KO lethality phenotype. However, whether the transgene fully rescued lethality or exhibited a fitness cost remained unclear. To address this question, we crossed females and males that encode for one copy of the transgene and quantified the progeny genotypes (Figure 3B). This scenario should lead to 50% of females having two copies of the transgene and 50% of males with one copy if there is no fitness cost. Indeed, we found that crossing both *Arp2D*- and *Arp2D2*-expressing flies led to at least 50% of the progeny being homozygous (female) or hemizygous (male) for the transgenes (p<0.05, Figure 3B). The slight increase over the expected 50% suggests that the X-chromosome balancer (*FM7*) has a small fitness cost compared to the transgenes. We conclude that when canonical Arp2 is replaced with either Arp2D or Arp2D2, no loss of fitness is detected under laboratory conditions (Figure 3B). We further tested whether a fitness cost would become apparent under environmental stress. We conducted the same cross with the most divergent duplicate (Arp2D) under heat stress at 29°C and found complete rescue, suggesting no fitness cost in viability even under these conditions (Figure S2A). We also did not notice any lifetime defects, abnormalities in morphology, or unusual behavior at room temperature or under heat stress. We conclude that the diverged Arp2D and Arp2D2 duplicates can functionally replace Arp2 for all assayed somatic roles.

### Arp2D-expressing males are subfertile

Having established that *Arp2D* and *Arp2D2* possess all the canonical *Arp2* functions, we next tested if they could fully complement *Arp2* functions in the male germline by comparing fertility of the ‘gene-replacement’ males. For this, we crossed hemizygous males with wildtype (Oregon R) females and, after one week of mating, compared adult progeny count to males encoding intron-less canonical *Arp2* in the *KO* locus (Figure 3C). We observed that *Arp2D*-expressing flies yielded fewer progeny than canonical *Arp2* (albeit this difference was not significant), whereas *Arp2D2*-expressing flies had progeny counts comparable to canonical *Arp2* (Figure 3C). Since males generally produce many sperm, one week of mating at room temperature may not readily reveal decreases in sperm count. Heat stress often exacerbates existing fertility defects, which become more detectable. We conducted the same male fertility assay as before, except at 29°C. We found that Arp2D-expressing males exhibited a significant decrease in adult progeny (p=0.02, Figure 3D), indicating they are subfertile. We then directly assessed sperm production by measuring the size of the seminal vesicle, where mature sperm are stored^29^. The area of the seminal vesicle scales with the number of stored mature sperm^29^. We found that the average area of seminal vesicles from *Arp2D*-expressing males was significantly smaller than those from canonical Arp2-expressing males, whether they mated or not (p=0.009, Figure 3E; p=0.003, Figure S2B). Therefore, *Arp2D*-expressing males produce less sperm and exhibit a fertility cost. We investigated whether the decrease in germline function was due to a lack of *Arp2D* expression in the testis and found that *Arp2D* and *Arp2D2* are both expressed in the testis at comparable levels (Figure S2C). Given *Arp2D* mRNA expression and the localization of Arp2D-GFP to testis actin (Figure 2C), we conclude that expression levels cannot account for the observed male subfertility. We conclude that Arp2D may be able to perform somatic roles in *D. melanogaster*, yet it lacks compatibility with *D. melanogaster* sperm development.

We also tested the fertility of *Arp2D*-expressing females. We crossed homozygous females to wildtype males and found a significant reduction in adult progeny (p<0.001, Figure S3A). We next investigated whether this is due to a lack of expression of Arp2D in ovaries. We imaged ovaries from wildtype *D. melanogaster* flies expressing GFP-tagged Arp2D and detected no Arp2D protein expression (Figure S3B). In contrast, we found that *D. pseudoobscura* canonical Arp2 does localize to actin-rich areas like the cortex (Figure S3B), suggesting that lack of Arp2D expression in the ovary is likely not due to a species difference in transcriptional or translational regulation. *D. auraria* Arp2D2-GFP was also expressed and localized to actin (Figure S3B). To further rule out the possibility that species-specific codon usage may lead to a lack of Arp2D expression in the ovary, we codon optimized *Arp2D* for *D. melanogaster* and inserted it in the *Arp2-*KO locus as done previously (Figure 3A). Because codon bias can also impact transcription^30^, we assessed mRNA expression and found that *Arp2D* mRNA remained the same as non-codon optimized Arp2D (Figure S3C), and female fertility remained extremely low (p<0.001, Figure S3D), suggesting subfertility is not due to species-specific codon usage. Given that Arp2D-GFP is expressed and localizes to actin in the testis, we conclude that reduced male fertility is due to Arp2D protein function, whereas failure to even express Arp2D protein in the ovary prevented us from further assessing its female germline function. Because Arp2D is natively expressed in the testis, we focused on investigating Arp2D protein function in the male germline.

### Arp2D-expressing males exhibit testis defects

We next investigated whether the testis exhibits any cytological defects that may lead to reduced sperm production. Sperm development can be tracked spatially in the fly testis (Figure 4A). Germline stem cells are at the apical end, where they differentiate and progress through mitosis, meiosis, and sperm maturation. After sperm elongation, sperm are separated by a process known as individualization, which is powered by actin cones. Fly testes appear morphologically as tubes coiled at the basal end, where individualizing sperm are found (Figure 4A). We were surprised that *Arp2D*-expressing testes exhibited a striking morphological defect in testis morphology (Figure 4B). The apical end of all *Arp2D*-expressing testes appeared swollen relative to the rest of the testis (p<0.001, Figure 4B). Because Arp2D was not codon-optimized for *D. melanogaster*, we considered this cytological defect might result from transcriptional and translational differences. However, codon-optimized *Arp2D*-expressing males exhibited the same abnormal morphology (Figure S4A). Alternatively, we hypothesized that this defect could be due to an artifact of *Arp2D* overexpression. Thus, we dissected the testes of male flies expressing two copies of wildtype *Arp2* and GFP-tagged *Arp2D* or *D. pseudoobscura Arp2* (four copies total). All exhibited wildtype morphology, suggesting the morphological defect is not due to overexpression of *Arp2D*, nor does *Arp2D* cause a dominant-negative effect (Figure S4B).

**Figure 4:**
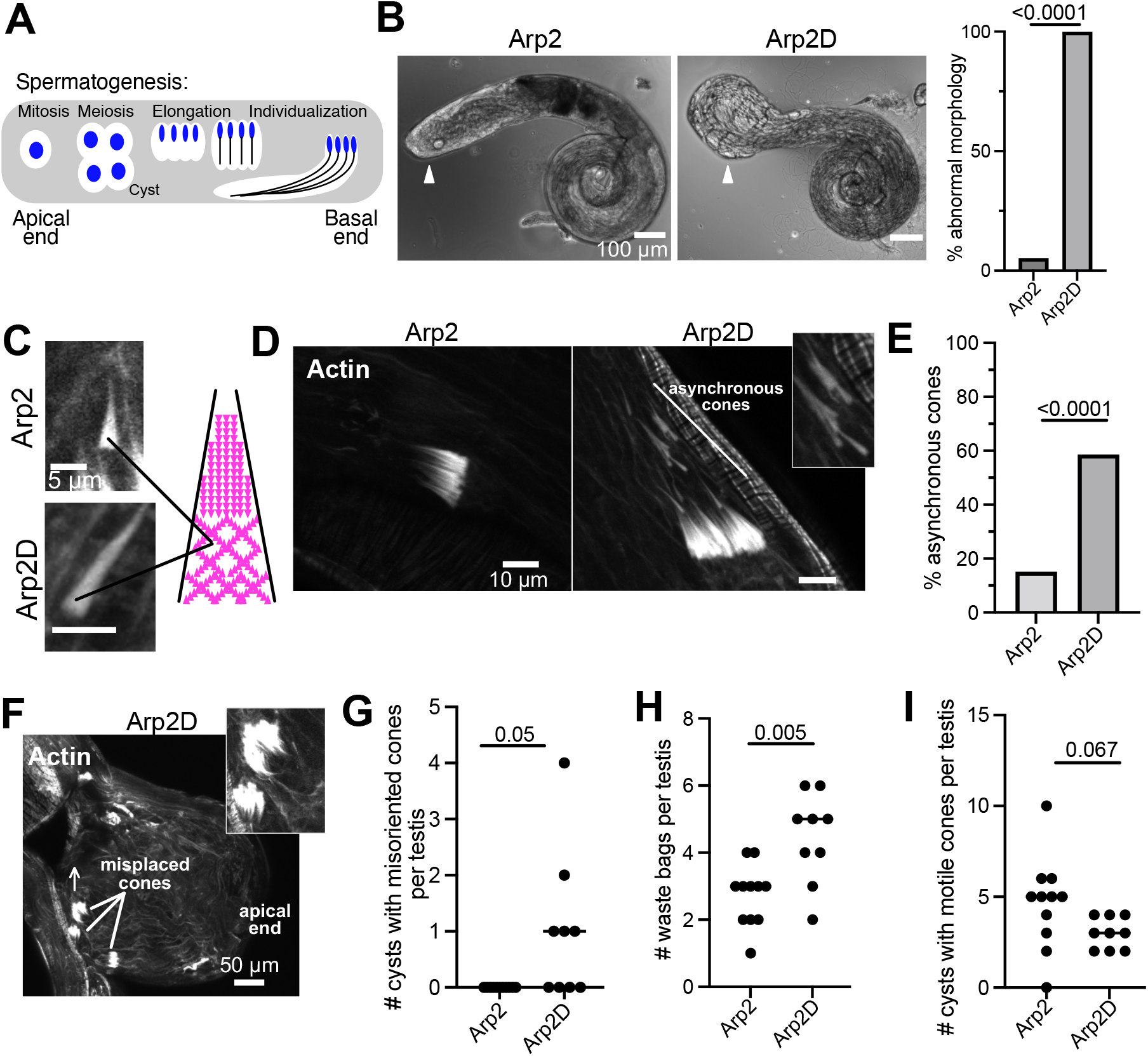
Arp2D-expressing males exhibit defects in sperm individualization. **A)** Schematic showing the stages of sperm development, including mitosis, meiosis, elongation, and individualization progressing from the apical end to the basal end. Nuclei are depicted in blue. **B)** Left, light microscopy image of testes from Arp2- and Arp2D-expressing flies. Arrowheads indicate apical end. Right, Arp2D-expressing flies exhibit an enlarged apical end, which was quantified as percent abnormal (Arp2: n=19; Arp2D: n=12). **C)** Images of a single actin cone containing canonical Arp2 or Arp2D. Cones are fan-like, indicating the presence of branched actin (depicted in cartoon). **D)** Images of dissected testes fixed and stained for actin. Testes expressing canonical Arp2 moved synchronously while cones in Arp2D-expressing flies frequently moved asynchronously. **E)** Quantification of cysts containing asynchronous cones (Arp2: n=21, Arp2D: n=14 testes). Arp2D-expressing flies exhibited significantly more asynchronous cones. **F)** Image of the apical end of Arp2D-expressing testes stained with actin. Motile cones are expected to face the apical end (the destination), yet some are misoriented and face the basal end. Arrow indicates direction cone front is facing. **G)** Quantification of cysts with cone fronts facing the basal end. None were seen in Arp2-expressing flies, whereas a significant number were seen in Arp2D-expressing flies. **H)** Quantification of the number of waste bags per testis. Arp2D-expressing testes exhibited more waste bags, which were identified by degraded actin cones. **I)** Quantification of cysts with motile cones, which are no longer localized to mature sperm nuclei. The number of cysts with motile cones per testis was reduced but insignificant for Arp2D-expressing flies.

The enlarged apical end of Arp2D-expressing testes is reminiscent of phenotypes previously observed in mutants in sperm individualization, which lead to an accumulation of debris and disorganized cysts with scattered actin cones^31-35^. Therefore, we investigated *Arp2D*-expressing testes for defects in individualization. We fixed and probed testes for DNA and actin to visualize actin cones in near-mature sperm. Actin cones in Arp2D-expressing males exhibited the normal fan shape that indicates the presence of the branched actin network near the cone’s front half^12^ (Figure 4C). However, we found that cones often moved asynchronously in Arp2D-expressing males, unlike canonical Arp2-expressing cones (p<0.0001, Figure 4D-E) or wildtype males^12^. Cones were also frequently misoriented. Cones normally move away from the basal end with the branched front facing the apical end, yet we often found the branched front of cones incorrectly facing the basal end (p=0.05, Figure 4F-G). We next assessed the last step of individualization: formation of the waste bag. Once actin cones reach the apical end, the excess cytoplasm and actin cones are degraded^10^. We quantified the number of waste bags (identified by actin cone degradation) and found that the number of waste bags per testis was significantly increased (p=0.005, Figure 4H). The increase in waste bags and the abnormal apical end was not due to a lack of caspase activity (Figure S4C), which is required for proper individualization^33,36,37^. We also found that the average number of cysts with motile cones per testis was slightly reduced, though not significantly (Figure 4I). Overall, defects were found throughout individualization, indicating Arp2D fails to carry out sperm individualization optimally in *D. melanogaster*.

### Arp2D’s divergent subdomain 2 loop contributes to testis defects

We next asked which structural divergence found in Arp2D leads to this testis phenotype. We compared the sequences of Arp2D orthologs to canonical Arp2 and Arp2D2 orthologs and found the most pronounced divergence in two segments: subdomain 2 and the C-terminus (Figure S5A). Canonical *Arp2* has a highly conserved alternately spliced exon in subdomain 2 (Figure S5A). Arp2D2 orthologs’ subdomain 2 appeared similar in length and sequence to the longer canonical *Arp2* splice variant, whereas *Arp2D* orthologs do not encode the alternate exon and notably diverge in sequence (Figure S5A). Most Arp2D2 orthologs have C-termini similar to canonical Arp2’s, yet Arp2D orthologs have significantly longer and diverged C-termini ranging from 10 to 21 additional highly charged amino acids, composed mainly of Lys (5-6 residues) and Asp/Glu (1-4 residues) (Figure S5A). Given canonical *Arp2*’s stringent conservation across eukaryotes, the sequence divergence of Arp2D suggests these regions may result in functional divergence.

We first tested if Arp2D’s divergent subdomain 2 loop contributes to the testis defects. We generated a chimera that replaces the Arp2D loop with the loop from *D. auraria* Arp2D2 (Figure 5A, Fig. S5B), which does not exhibit reduced fertility or an abnormal testis morphology. The chimera’s expression was comparable to that of Arp2D at the mRNA level (Figure S6G), and hemizygous males were viable like Arp2D-expressing flies. We found that testes were strikingly normal in morphology (Figure 5B). While 100% of Arp2D-expressing testes were bulbous, only 30% of the chimera’s testes exhibited a bulbous apical end (p<0.001, Figure 5C). However, the phenotypic rescue was incomplete, as cones remained asynchronous (Figure 5D, Figure S6A-E). We conclude that the subdomain 2 loop contributes to Arp2D’s deficient germline function, yet it is not the only structural divergence leading to the observed defects.

**Figure 5:**
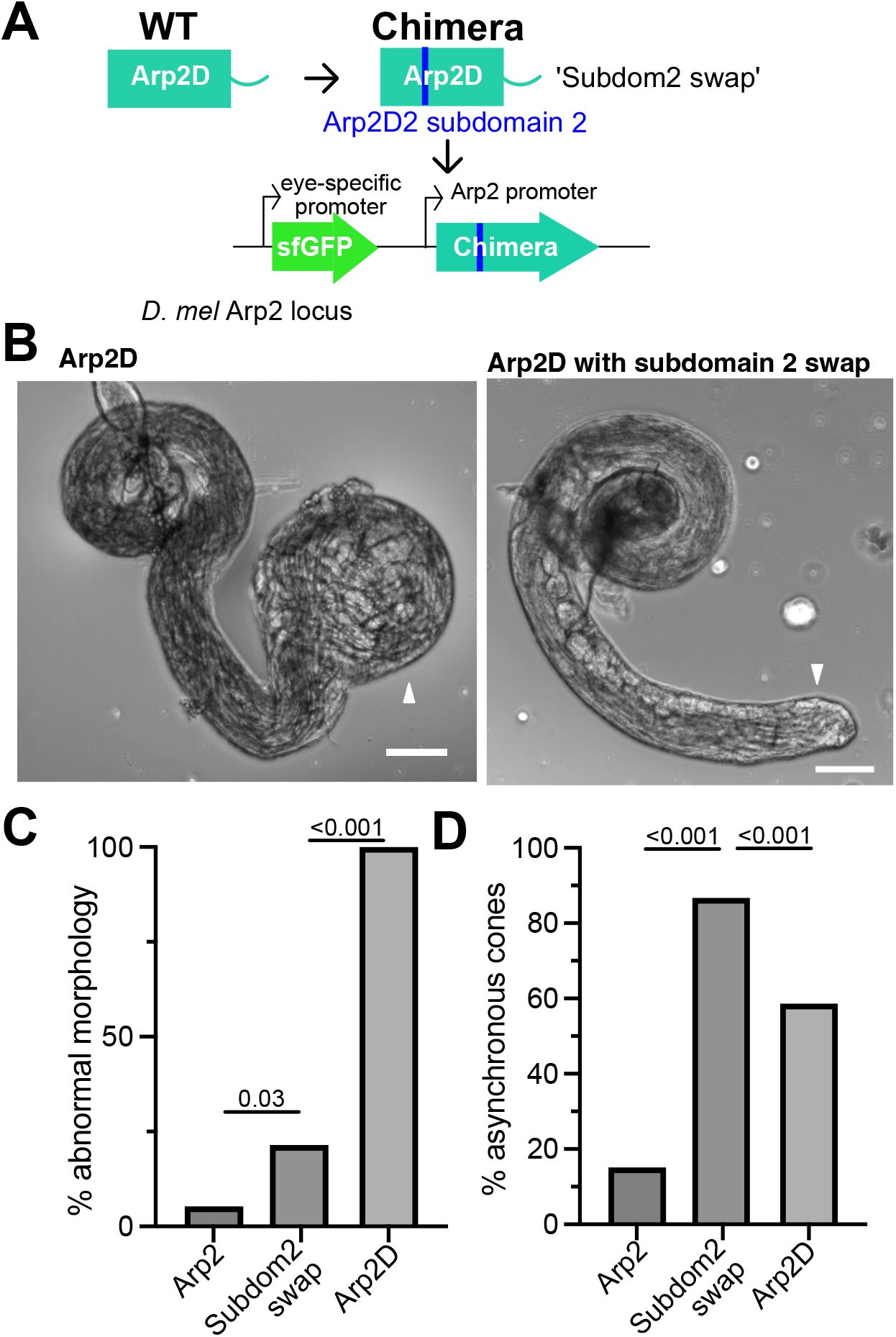
Arp2D’s divergent subdomain 2 loop contributes to testis defects. **A)** Schematic showing subdomain 2 swap. *D. melanogaster* Arp2-KO flies were genetically modified as in Figure 3A except *Arp2D* (*D. pseudoobscura*) encoded subdomain 2 from *Arp2D2* (*D. auraria*). **B)** Brightfield images of one-day old virgins expressing Arp2D or ‘subdom2 swap’ in (A). Arp2D’s apical end is an abnormal bulbous shape while the chimera exhibited normal morphology at the apical end. Arrowheads indicate apical ends, and scale bars are 100 μm. **C)** Quantification of the percent of testes with an abnormal (bulbous) apical end (Subdomain 2 swap: n=28; Arp2 and Arp2D data from Figure 4B). **D)** Quantification of the percent of cysts containing asynchronous cones. (Arp2D2: n=12; data for Arp2 and Arp2D is the same as in Figure 4E. Despite a normal apical end, the chimera still exhibits asynchronous cones.

### Arp2D’s C-terminus is required for its activity in somatic tissue but reduces optimal function in the testis

We next tested if Arp2D’s unique C-terminus also affects sperm development. We shortened *D. pseudoobscura* Arp2D’s C-terminus by removing 11 amino acids, including the lysines conserved among most *obscura* species, and replaced it with Arp2’s C-terminus, which is only 3 residues long (Figure 6A, Figure S5C). We inserted the chimeric transgene (‘Arp2D-2CT’) into the *Arp2*-KO locus to test its impact on testis function (Figure 6A). Similar to previous designs, this transgene encoded canonical Arp2’s 5’UTR and 3’UTR (Figure 3A). Although we removed divergence in Arp2D’s C-terminus, Arp2D-2CT surprisingly failed to rescue *Arp2*-KO lethality (p<0.0001, Figure 6B). We verified that the construct was expressed and comparable to full-length Arp2D expression in heterozygotes (Figure S6F). Therefore, Arp2D’s evolutionarily unique C-terminus is critical for its activity.

**Figure 6:**
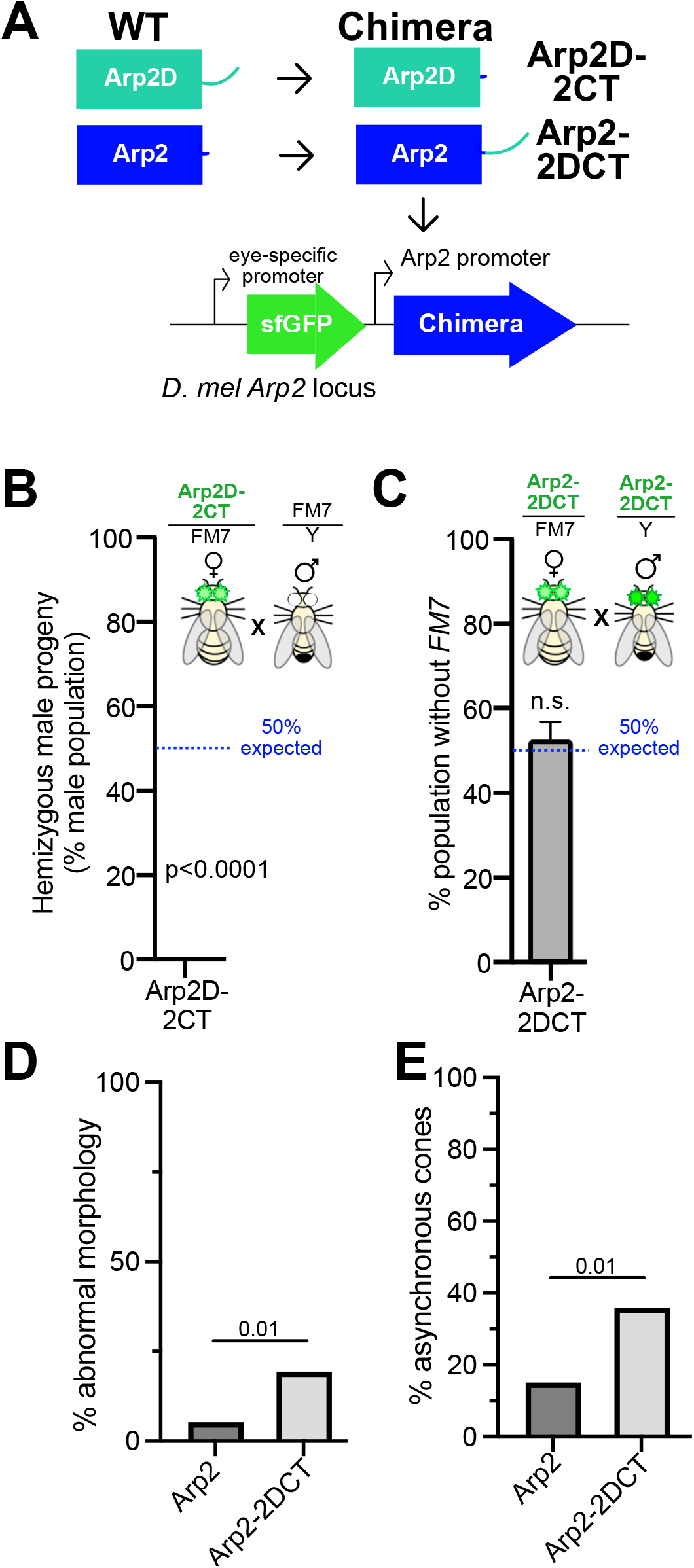
The unique C-terminus is required for Arp2D function but leads to testis defects in *D. melanogaster*. **A)** Schematic showing the generation of two chimeras. The extended C-terminus of Arp2D was replaced with Arp2’s short C-terminus to produce the ‘Arp2D-2CT’ chimera. The extended C-terminus of Arp2D was engineered on to canonical Arp2 to form the ‘Arp2-2DCT’ chimera. The transgenes were inserted into the *Arp2*-KO locus as done in Figure 3A. **B)** Schematic of the cross used to test for rescue of *Arp2*-KO lethality with chimeric Arp2D: *Arp2D-2CT*. Heterozygous *Arp2D-2CT*-expressing females were crossed to males with the balancer *FM7* (5 replicate crosses). Green eye color indicates presence of the transgene, with heterozygous females having dim GFP-positive eyes. Percent of the male progeny that was hemizygous for *Arp2D-2CT* is shown. No viable hemizygous males were produced (5 replicate crosses). **C)** Schematic of the cross used to test for rescue of *Arp2*-KO lethality with chimeric Arp2: *Arp2-2DCT*. Heterozygous *Arp2-2DCT*-expressing females were crossed to hemizygous males (6 replicate crosses). Quantification shows approximately 50% of the cross’s progeny expressed only *Arp2-2DCT* with no *FM7* present, indicating a complete rescue of *Arp2-*KO lethality. Genotype fractions in B and C were compared to Mendelian expectation using a chi-squared test. **D)** Percent of abnormal testes (bulbous apical end) is shown for Arp2- and Arp2-2DCT-expressing flies (Arp2-2DCT: n=31; data for Arp2 from Figure 4B). **E)** Percent of cysts containing asynchronous cones is shown for Arp2- and Arp2-2DCT-expressing flies (Arp2-2DCT: n=24; data for Arp2 from Figure 4E).

We next tested the impact of Arp2D’s C-terminus on canonical Arp2. We modified the *Arp2*-KO flies by inserting canonical Arp2 with Arp2D’s C-terminus (Figure 6A, Figure S5D). The transgene was expressed (Figure S6G), and the *Arp2*-KO lethality phenotype was fully rescued (Figure 6C). However, we found a significant increase in testes with an enlarged apical end (p=0.01, Figure 6D) and actin cones that moved asynchronously (p=0.01, Figure 6E). The number of waste bags per testis, number of motile cones, and orientation of cones were comparable to that of canonical Arp2 (Figure S6A, D-E). Therefore, despite the importance of the unique C-terminus for Arp2D function, this domain negatively impacts *D. melanogaster* sperm development.

## Discussion

Arp2 arose early in eukaryotic evolution and is under stringent sequence conservation for its many roles throughout all tissue types^19,38^. Our studies reveal recurrent Arp2 gene duplication and subsequent sequence diversification in *Drosophila*. The paralogs’ exclusive expression in the testis suggests a selective pressure to have a testis-specific Arp2. Their diversification may have retained Arp2 function, sub-functionalized, or led to an entirely new function that does not involve generating branched actin. We found that they can functionally replace Arp2 and lead to viable flies, even in a non-native species. However, Arp2D’s considerable divergence in sequence and specialization for male germline roles in the *obscura* clade appears to have led to incompatibility with *D. melanogaster* sperm development. This study reveals how the Arp2 structure can be altered and sub-functionalize Arp2.

Gene duplications often relieve tension between conflicting roles of the parental gene^39-42^. We find that Arp2 may have conflicting roles between its ‘housekeeping’ functions in somatic tissue and those in sperm development, and the optimal Arp2 sequence for somatic roles may result in a fitness cost in the germline. This tension can be relieved with a germline-specific Arp2 paralog. Surprisingly, evolution has selected for a testis-specific Arp2 paralog twice, suggesting a recurrent need for an Arp2 that is specialized for male germline roles. Arp2D2 and Arp2D’s sequence and functional differences suggest they have evolved uniquely for Arp2’s spermatogenic roles specific to their respective species. Although we found no fertility cost in *Arp2D2*-expressing *D. melanogaster* flies, Arp2D2 may have a subtle deviation from canonical Arp2’s function. Alternatively, Arp2D2 may function exactly like Arp2, and the *montium* species require a higher copy number in the testis. Intriguingly, Arp2 appears to be the only member of the Arp2/3 complex that has repeatedly duplicated. We see one additional instance of Arp2/3 diversification in *Drosophila*; *D. melanogaster* encodes a gene duplicate of Arpc3 exclusively expressed in the ovary and localizes to actin structures found only in the female germline^9^. The diversification of multiple *Drosophila* Arp2/3 subunits indicates that each member of the complex may differentially tune actin polymerization for specific germline roles, and Arp2 uniquely plays a central function in testis specialization.

Despite being only 70% identical to canonical Arp2, Arp2D surprisingly does not lead to a fitness cost in *D. melanogaster* viability, yet subtle defects in somatic tissues may exist and remain undetected. We focused our analysis on the striking testis phenotypes due to Arp2D’s male-specific expression in the *obscura* species^23^. Although Arp2D appears to generate branched actin networks in the *D. melanogaster* testis (Figure 4C), Arp2D fails to perform like canonical Arp2. The kinetics of complex assembly or Arp2D-catalyzed actin polymerization may be changed. Alternatively, the Arp2D/3 complex may lead to branched actin networks that are unstable or hyper-stable. There is precedence for diversification of Arp2/3 complexes leading to an impact on polymerization rate (gene duplicates of ArpC1 and ArpC5^20^) or network stability (gene duplicate of human Arp3^21^). These differences may not be attributed directly to the complex but may result from the loss or gain of interactions with regulators^19^. A testis-specific factor in the *obscura* species may be required for Arp2D to function normally in *D. melanogaster* testes.

We show that Arp2D’s structural divergence from canonical Arp2 in subdomain 2 contributes to the observed testis defects in *D. melanogaster*. Canonical Arp2’s subdomain 2 loop has limited structural information, partly due to its flexibility and lack of electron density in high-resolution structures^14,16^. However, we have a few clues about the functional impact of subdomain 2. In canonical Arps 2 and 3, it faces towards the mother actin filament^15,16,43^. One study showed Arp3’s subdomain 2 loop facilitates interaction with actin, and by removing it, the rate of actin polymerization increases, suggesting it is kinetically important^43^. Because our study shows Arp2D’s subdomain 2 is incompatible with *D. melanogaster* sperm development, this structural region carries functional importance and may enable critical interactions with actin that impact kinetics like Arp3’s loop. Interestingly, canonical Arp2’s subdomain 2 undergoes alternative splicing, resulting in two splice variants differing only by five amino acids^44,45^ (Figure S5A). Although nothing is known about the distinct functions of Arp2’s splice isoforms, we speculate that modulation of Arp2’s subdomain 2 can tune actin polymerization for different contexts, and perhaps Arp2D has evolved a divergent loop that specializes polymerization kinetics specifically for sperm development in *obscura* species.

We also find that Arp2D’s C-terminus is incompatible with *D. melanogaster* sperm development. Because canonical Arp2 can be C-terminally tagged without perturbing activity^28^ or localization (Figure 2C), the functional impact of Arp2D’s C-terminus on Arp2 likely depends on sequence and not length. Unlike subdomain 2, much less is known about the role of canonical Arp2’s C-terminus in actin polymerization and how altering its sequence may impact function. However, a major clue comes again from Arp3 studies. Arp3’s C-terminal tail serves an autoinhibitory role by binding the hydrophobic groove between subdomains 1 and 3, keeping the Arp2/3 complex in an inactive conformation until ATP^46^ or a nucleation-promoting factor (WASP)^47^ binds. Analogously, it would seem Arp2D’s C-terminus may be inhibitory, yet this domain is highly charged and unlikely to bind the hydrophobic groove of Arp2. Instead, Arp2D’s C-terminus may alter charge-charge interactions with regulators critical for actin polymerization in *D. melanogaster* individualization.

Despite its negative impact on sperm development, Arp2D’s unique C-terminus is surprisingly required for Arp2D’s somatic function. Because the C-terminus is predicted to be unstructured, removing it is unlikely to make the protein less stable and, therefore, must be required for Arp2D protein function. The N-terminal domain composing the majority of Arp2D appears structurally like canonical Arp2 based on homology predictions; thus, we expected this domain would be sufficient to interact with actin and the rest of the Arp2/3 complex for actin polymerization. However, because the unique C-terminus is required for somatic function, we infer that Arp2D’s C-terminus has co-evolved with the ‘Arp2-like’ N-terminal domain and is now integral for full-length protein activity, yet incompatible with the evolutionarily older canonical Arp2 sequence for testis function. The C-terminus may directly stimulate Arp2D for nucleation activity or have acquired a regulator binding site. Interestingly, Arp2D orthologs’ C-termini vary in length and overall charge (Figure S5A). We speculate that each ortholog’s C-terminus has co-evolved with its respective ‘Arp2-like’ domain, which has led to species-specialized germline function. Because the *obscura* species retain canonical Arp2, Arp2D had the opportunity to evolutionarily explore new sequences and roles in sperm development without compromising viability, as canonical Arp2 preserved somatic function.

The question remains why Arp2 has recurrently specialized exclusively for testis function. Sperm development is under intense selective pressure from sperm competition and sexual selection^48,49^. Consequently, biological innovation frequently occurs in the testis, leading to significant species variation in sperm developmental processes that likely have unique cytoskeletal requirements. For example, actin cones, found only in the *Drosophila* male germline, arose for the syncytial nature of insect germ cell development^10^, and their construction and motility may require a specialized Arp2/3-generated branched actin network. Our findings show that by studying how Arp2 diversification impacts testis actin, we can gain insight into how Arp2 adapts structurally and functionally for the germline.

## Materials and Methods

### Culturing flies and sequencing the *Arp2D2* locus

*Arp2D2* was identified by using a tBLASTn search with canonical *D. melanogaster* Arp2, and mRNA encoding an ‘Arp2-like’ gene was found in *D. serrata* (Accession XM_020944101.1) and *D. kikkawai* (Accession XM_017179971.1). The coding sequence was then mapped to the *D. kikkawai*^*24*^ and *D. serrata*^*25*^ genomes (contig 7911 and MTTC01000122.1, respectively) to identify the loci and found they were syntenic and differed in canonical Arp2’s locus. For sequencing the shared syntenic locus of *montium* species, genomic DNA was obtained from 10-15 whole flies as done previously^23^. *D. melanogaster* and *D. kikkawai* genomes were aligned, and primers were designed to target conservation in the intergenic regions flanking the *D. kikkawai Arp2D2* locus. Primers were then used for targeted sequencing of the *Arp2D2* locus in unsequenced *montium* species. Some *montium* species had genomic DNA purified for a past study^50^ and were kindly offered for the study. Cultured species for which genomic DNA was not already available were homogenized and DNA was extracted as done previously^50^. PCRs were conducted following a touchdown protocol^51^ with Phusion, following the manufacturer’s instructions (NEB). PCRs were directly sequenced, or first TOPO cloned (Invitrogen) followed by sequencing. Primers were iteratively designed using sequences from successful PCRs to obtain *Arp2D2* loci sequences that were more diverged in the intergenic regions. All flies were cultured on yeast-cornmeal-molasses-malt extract medium at room temperature. Cultured species and primer sequences are listed in Data S1, and *Arp2D2* sequences obtained by PCR are provided in Data S2.

### Identification of *Arp2* and *Arp2D2* in sequenced *montium* genomes

*D. melanogaster* Arp2 and *D. kikkawai* Arp2D2 protein sequences were used in a tBLASTn search to obtain orthologs in sequenced *montium* species^26^. The search set was BioProject ID 554346 (a whole genome shotgun contigs database)^26^. Hits that had E values of 0 were obtained and syntenic orthologs of *D. kikkawai Arp2D2* were identified by conservation of upstream and downstream genetic regions including neighbors *Hmu* and *CG3339*. To obtain syntenic *Arp2* orthologs, upstream and downstream regions including genetic neighbors *Pp2B-14D* and *CG9903* were identified. *Arp2D2* coding sequences were then extracted based on alignments with *D. kikkawai Arp2D2*, and *Arp2* coding sequences were obtained by aligning with *D. kikkawai Arp2* or *Arp2* from a more closely related *montium* species. All analyses were done using the Geneious software package^52^. Accession numbers of scaffolds encoding *Arp2D2* or *Arp2* in BioProject 554346^26^ are provided in Data S1.

### Phylogenetics and structural analysis

Nucleotide sequences were aligned using translation alignment in the Geneious software^52^. A maximum-likelihood tree (Data S3) was generated with PhyML (HKY85 substitution model), and 100 bootstrap replicates were performed for statistical support in Geneious^52^. Protein sequence alignments were conducted using MUSCLE^53^ in Geneious^52^. For structural analyses, a homology model of *D. pseudoobscura* Arp2 was obtained using SWISS-MODEL^54^, which used PDB 4JD2^55^ as a template, and structures were viewed using Chimera^56^.

### Generation of fly transgenics

#### Generation of GFP-tagged Arp2 and paralogs

To visualize Arp2 variants, *D. pseudoobscura Arp2* (intronless, short splice variant), *D. pseudoobscura* Arp2D, and *D. auraria Arp2D2* were C-terminally tagged with superfolder GFP and cloned into a vector encoding *attB*. Each construct was cloned with either the promoter for *D. melanogaster Arp2* or for *D. melanogaster Arp53D* (a gene highly expressed in the testis). Constructs were midi-prepped (Takara Bio), and for site-directed transgenesis, constructs were co-injected with PhiC31 into a fly line encoding *attP* on the third chromosome (stock 9744 from Bloomington *Drosophila* Stock Center) by Rainbow Transgenic Flies, Inc.

#### Generation of Arp2 Knockout

Arp2 was knocked out using CRISPR/Cas9 and replaced with *DsRed* to track the knockout allele. The guide RNAs were chosen based on a low probability of off-targets (http://flyrnai.org/crispr2/) and targeted in the intergenic regions upstream and downstream of the coding regions to completely remove the gene. The guide RNAs (GTAGCTGCTACTAGCAGACT and TCGTTACTCCCCAGAGTTGA) were cloned into pCFD4 (Addgene plasmid 49411)^57^ and the 1kb regions upstream and downstream of the cut sites were cloned into vector pHD-attP-DsRed (Addgene plasmid 51019)^58^. Plasmids were midi-prepped (Takara Bio), and BestGene, Inc. injected them into a nanos-Cas9 expressing fly line (51324 from Bloomington *Drosophila* Stock Center). G0 adults were crossed to w1118 flies to identify transformants (DsRed-fluorescent eyes) and cross out Cas9. Transformants’ X-chromosome was balanced with *FM7*. We sequence verified CRISPR/Cas9 cut sites and presence of *DsRed*. The KO allele was only found in heterozygous females, as expected for the *Arp2*-KO phenotype, and no homozygous or DsRed-positive males were identified.

#### Generation of transgenic lines with Arp2-KO flies

The genes encoding *D. pseudoobscura Arp2D* and *D. auraria Arp2D2* were obtained by PCR from the species’ genomic DNA. Canonical Arp2 was obtained by PCR from *D. melanogaster* cDNA, and the short splice variant was used for this study. Codon-optimized Arp2D was synthesized by Integrated DNA Technologies, Inc. All mutants (Arp2D subdomain 2 swap, Arp2D-2CT, and Arp2-2DCT) were generated using Q5 site-directed mutagenesis (NEB). All genes were cloned into a vector encoding an attB site and superfolder GFP under the control of an eye-specific promoter (3XP3). Flanking all genes were approximately the 1kb upstream and downstream regions of canonical *D. melanogaster* Arp2. Constructs were midi-prepped and injected into the *attP*-encoding *Arp2* KOs by Rainbow Transgenic Flies, Inc. We crossed G0 adults to *Arp2*-KO stocks, selected flies with GFP- and DsRed-positive eyes, and balanced the X-chromosome with *FM7*. To sequence verify all transgenics, genomic DNA from flies was obtained by grinding two flies in 10 mM Tris-HCl pH8, 1 mM EDTA, 25 mM NaCl, and 200 μg/mL Proteinase K. The lysate was incubated at 37°C for 30 minutes then at 95°C for 3 minutes to deactivate Proteinase K. The lysate was briefly centrifuged, and the supernatant was used for PCR-amplification of the modified loci. A touchdown protocol^51^ was conducted with Phusion (NEB), and PCR products were used for Sanger sequencing. Sequences for all constructs used to generate transgenic fly lines are provided in Data S4.

### RT-PCR

To detect expression in the germline, abdomens were separated from the body and further dissected in PBS to extract testes or ovaries. Tissues or whole flies (5-10 total) were homogenized with TRIzol (Invitrogen) then briefly centrifuged to separate the supernatant. The supernatant was chloroform-extracted, resulting in a soluble phase that was then isopropanol-extracted. The RNA that precipitated was briefly centrifuged, and the resulting pellet was washed with 70% ethanol and resuspended in RNAse-free water. Samples were treated with DNase I (Zymo Research), followed by purification and concentration (RNA Cleanup & Concentrator kit, Zymo Research). The samples were used to generate cDNA using SuperScript III First-Strand Synthesis, following the manufacturer’s instructions (Invitrogen). The cDNA was then used to conduct RT-PCRs with Phusion (NEB). Primers are provided in Data S1.

### Imaging

#### Ovary live imaging

Virgin females were fed yeast paste (Sigma Aldrich) for two days prior to ovary dissections. Ovaries were dissected in PBS then incubated with Hoechst 33342 (Invitrogen) for 10 minutes at room temperature and washed once with PBS. Images were then collected on Nikon CSU-W1 with SoRa (UTSW Live Imaging Core).

#### Testis imaging

For brightfield imaging of testes, similarly aged males were collected and dissected in PBS. They were then placed on slides for live imaging. Images were collected on a Nikon Ts2 epi-fluorescent microscope with a 10x objective.

For confocal imaging, virgin male flies were dissected at 0-3 days old. For live imaging of GFP-tagged proteins at cones, testes were dissected in PBS and then transferred to a slide and torn to separate cysts. Cysts were then incubated with 10 μM SiR-Actin^59^ (Cytoskeleton, Inc.) and Hoechst 33342 (Invitrogen) for 5-10 min before imaging. For fixed whole mount testes and seminal vesicles, germline tissue was dissected in PBS and then transferred to an eppendorf tube for fixation using 4% PFA (25 min at room temperature). Samples were then washed twice for 15 min each with PBS including 0.1% Triton. To probe for actin and DNA, testes were stained with 2 μM SiR-Actin^59^ (Cytoskeleton, Inc.) for 2-3 hours at room temperature and incubated with Hoechst 33342 (Invitrogen) for 10 min at room temperature. Stained testes were then placed on a slide, non-germline tissue was further removed, and a coverslip was placed following addition of ProLong Diamond Antifade Mountant (Thermo Fisher). To probe for activated caspase 3, whole mount testis imaging was conducted similarly, though testes were stained with anti-cleaved caspase 3 (Asp 175, Cell Signaling) as done previously^36^ and probed with anti-rabbit secondary antibody (Alexa Fluor 488, Invitrogen). All testis imaging was done using an inverted Ti2 Nikon AX-R confocal microscope and NIS-Elements (Nikon).

### Quantification of male germline defects

Z-stacks were acquired of whole mount testes that were stained for actin and DNA. Stacks were then visualized with NIS-Elements (Nikon) to quantify testis defects, including motile cones, asynchronous cones, misoriented cones, and waste bags. Cones that were no longer localized to sperm nuclei were classified as ‘motile.’ Motile cones were quantified as asynchronous if a cyst had more than two cones that were trailing behind most cones. Waste bags were identified by degrading, amorphous actin cones. Misoriented cones were identified by the front of actin cones facing the basal end instead of the apical end. When counts of cysts or waste bags were compared, the statistical significance was evaluated with a t-test. When the percentage of asynchronous cones and testes with abnormal morphology were compared, statistical significance was assessed using a chi-squared test, using values for Arp2-expressing flies as the expectation. Seminal vesicle size was measured using the line tool in Image J^60^, and a t-test was used to evaluate statistical significance.

### Fly crosses

Oregon-R flies were used for wildtype flies. For all crosses, female virgins that were 1-5 days old and male virgins that were 1-3 days old were used. Virgins were maintained at room temperature until crosses were set up. All crosses were set up with females in excess of males at a ratio of 5:2. Pairs were mated for approximately one week with vials being flipped every 3 days. For heat stress, the mating pairs were maintained at 29°C, and all crosses were carried out with consistent light/dark cycles. Adult progeny were quantified for no more than 16 days relative to when crosses were setup or 13 days (crosses at 29°C) to avoid counting progeny from the next generation. Parental flies that died during the mating week were tallied and did not differ significantly among genotypes. All fertility assays were done at least three times, and statistical significance was evaluated with a t-test. For assays testing for rescue of *Arp2*-KO lethality, at least 5 replicates were done (noted in legends), and genotypes were scored per vial. Averages with variance are displayed. To assess statistical significance, the genotypes were summed across replicates and the observed ratio of genotypes was compared to the expected Mendelian proportion with a chi-squared test. A p-value less than 0.05 resulted if observed ratios deviated significantly from the Mendelian expectation.

## Acknowledgments

We thank Michael Buszczak, Lisa Kursel, and Harmit Malik for their comments on the manuscript and helpful discussions. We also thank Lisa for genomic DNA of the *montium* species. We thank Dean Smith, Elizabeth Chen, and John Abrams for experimental suggestions, the CC3-antibody (Abrams lab), and a homogenizer (Smith lab). Many *Drosophila* species were obtained from the National *Drosophila* Species Stock Center (Cornell University), and transgenic flies were generated by Rainbow Transgenic Flies, Inc. We thank the Sanger Sequencing Cores at Fred Hutchinson Cancer Center and UT Southwestern, as well as UT Southwestern’s Quantitative Light Microscopy Core, a Shared Resource of the Harold C. Simmons Cancer Center, supported in part by an NCI Cancer Center Support Grant, 1P30 CA142543-01. This work was initiated using an NIGMS K99 Pathway to Independence Award GM137038 (CS) and funds to Harmit Malik from NIGMS grant R01GM074108 and the Howard Hughes Medical Institute. Work is currently funded by a recruitment award from the Cancer Prevention and Research Institute of Texas (CS), the UT Southwestern Endowed Scholars Program (CS), a UT Southwestern STARs Fund (CS), and an NIGMS R00 (GM137038, CS).

## Supplementary Figures

**Figure S1:**
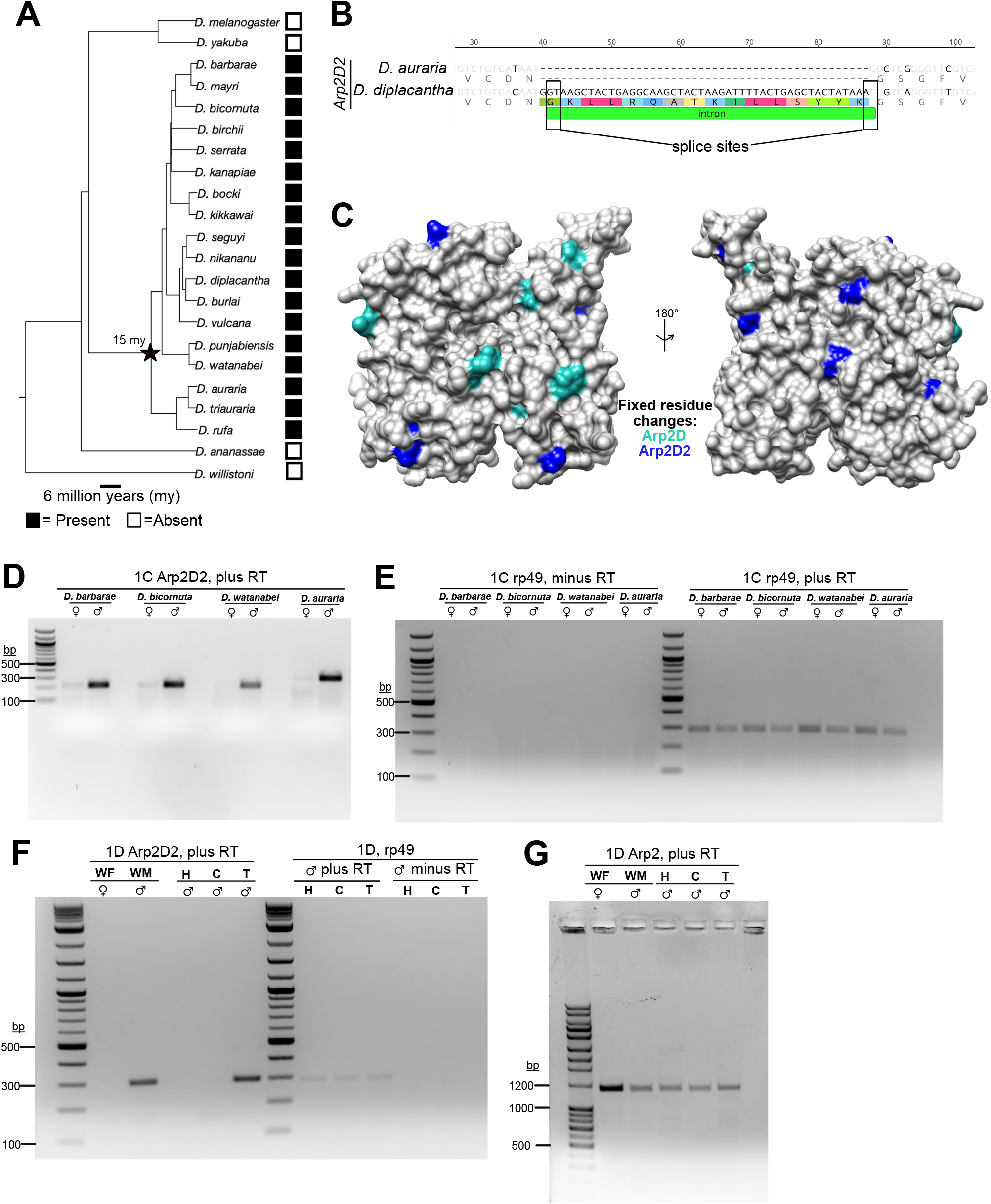
Evolutionary analyses and controls for RT-PCRs in Figure 1. A) A *Drosophila* species tree including 18 *montium* species. The *Arp2D2* syntenic locus was PCR amplified and sequenced. Presence and absence of *Arp2D2* in each species is indicated with black and white boxes, respectively. **B)** An alignment of *D. auraria* and *D. diplacantha Arp2D2* is shown, and the single intron and splice sites are annotated in *D. diplacantha Arp2D2*. **C)** Space filling representation of a *D. pseudoobscura* Arp2 homology model displaying residues that are uniquely conserved in either Arp2D or Arp2D2. Conserved Arp2D residues are shown in teal and Arp2D2 conserved residues are blue. No conserved residue changes were shared between Arp2D and Arp2D2. **D)** The full gel in Figure 1C’s top image, showing RT-PCRs of *Arp2D2*. **E)** The bottom image of Figure 1C showing RT-PCRs of *Arp2D2*. Template in the left side of the gel was cDNA synthesis conducted without reverse transcriptase (RT), and lack of signal indicates no genomic DNA contamination. Template on the right side of the gel was cDNA synthesis done with RT. Signal for *rp49* is comparable among samples. **F)** The full gel for the top image in Figure 1D. The left side of the gel corresponds to the top image (plus RT), and the right side includes RT-PCRs with the same samples except *rp49* expression is shown. The minus RT lanes indicate no genomic DNA contamination. **G)** The full gel for the bottom image in Figure 1D.

**Figure S2:**
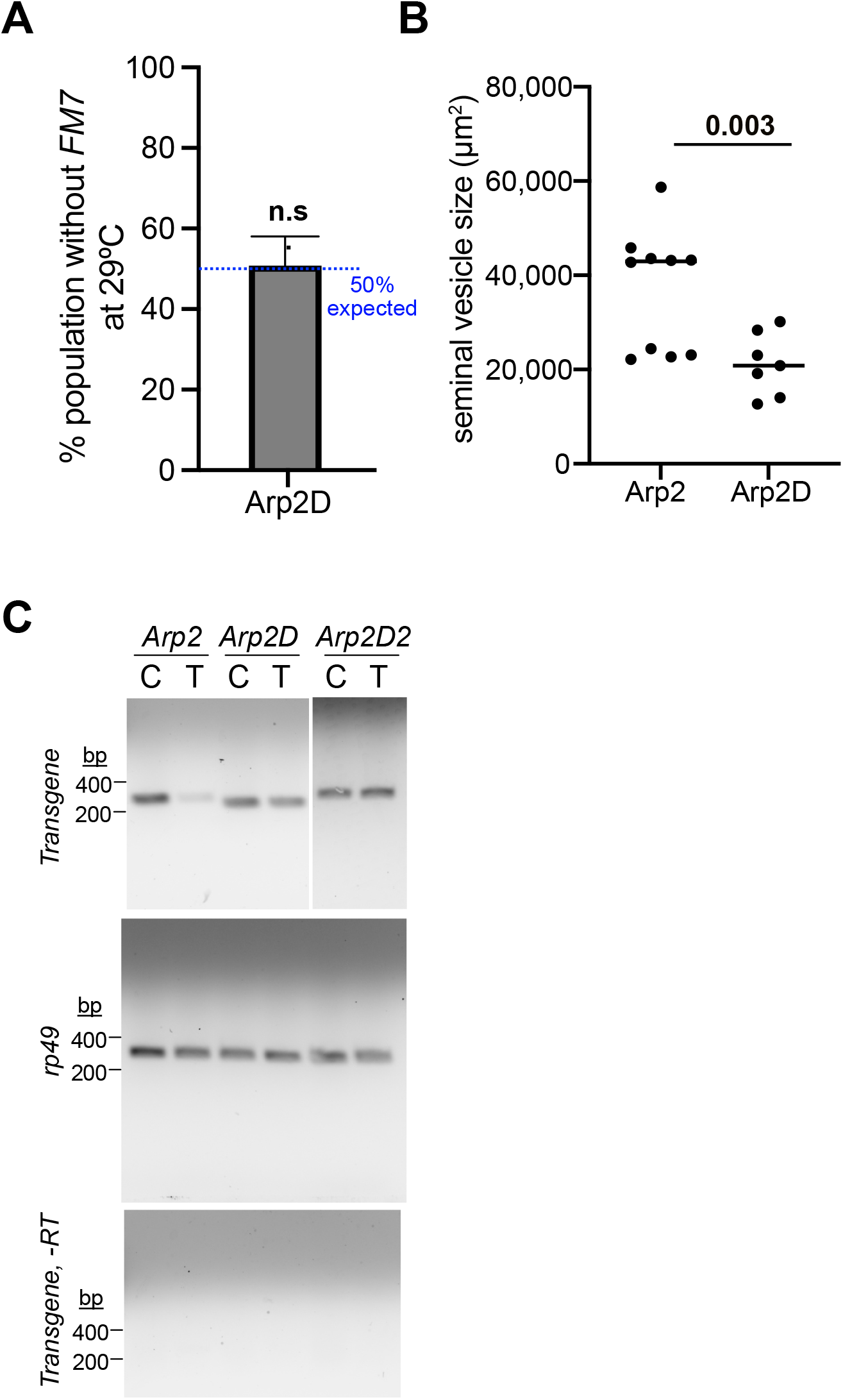
Tests for fitness cost and expression of Arp2D-encoding flies. **A)** The same cross with Arp2D-expressing flies in Figure 3B was conducted at 29°C to test for rescue under heat stress (7 replicate crosses). Percent of the progeny population that had no *FM7* (*Arp2D*-expressing homozygous females and hemizygous males) is shown and was not significantly different from the expected percent based on Mendelian inheritance (50%), indicating a full rescue. **B)** Quantification of seminal vesicle area from similarly aged Arp2D-expressing virgin males (3 d old) consistently exhibited significantly smaller seminal vesicles. **C)** RT-PCRs assessing expression of canonical *Arp2, Arp2D*, and *Arp2D2* in the fly lines depicted in Figure 3A. Male flies were dissected, separating carcass (‘C’) and testis (‘T’). *rp49* is shown to compare overall cDNA levels of the samples. The bottom gel image indicates cDNA samples generated without reverse transcriptase. No genomic DNA contamination was detected.

**Figure S3:**
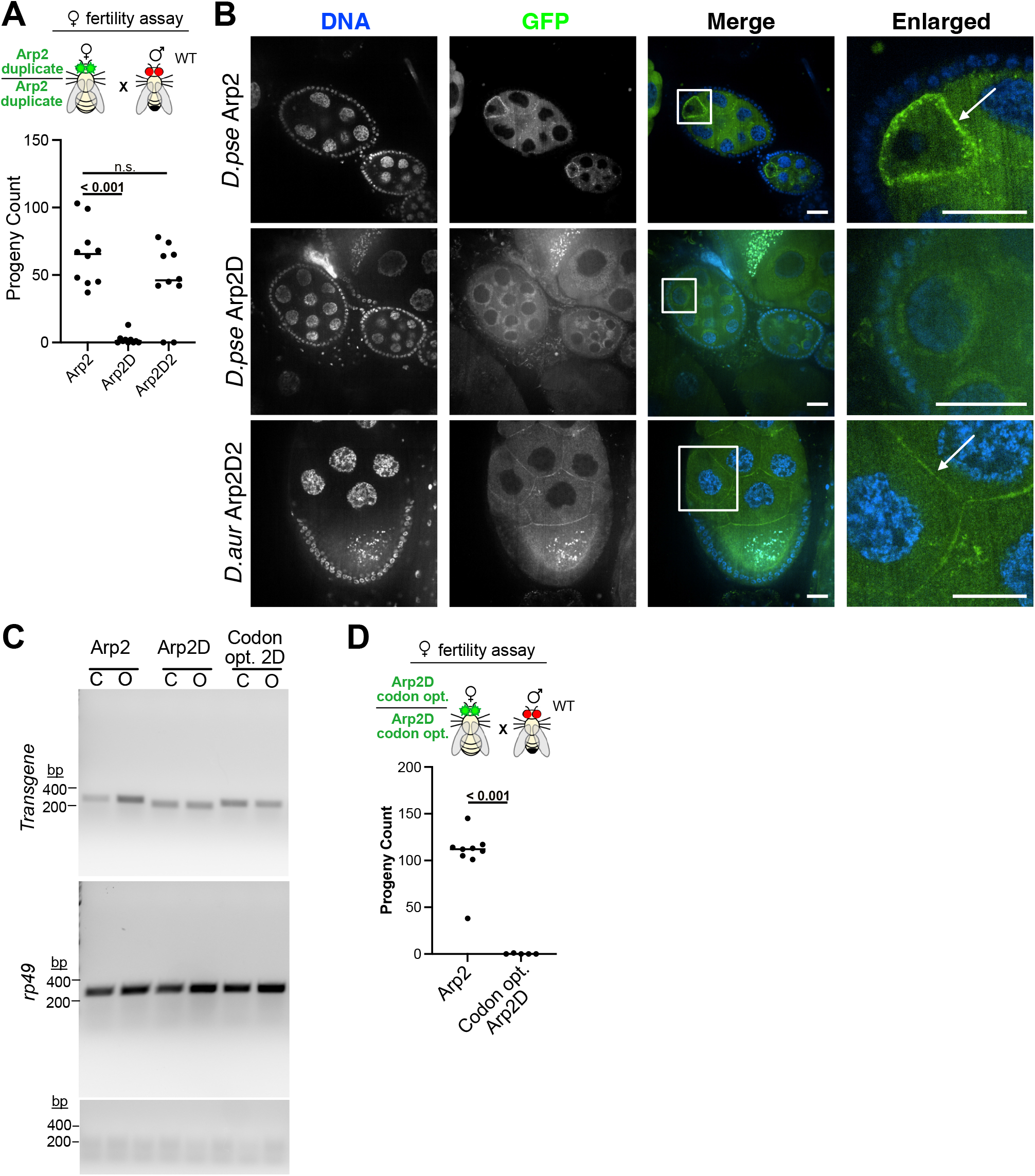
*D. pseudoobscura Arp2D* protein is not expressed in *D. melanogaster* ovaries unlike testes. **A)** Schematic for a female fertility assay. Arp2-, Arp2D-, or Arp2D2-expressing females were crossed to Oregon R (wildtype) males. Total adult progeny is shown, indicating Arp2D-expressing females were significantly subfertile. **B)** Live imaging of ovaries from *D. melanogaster* females expressing GFP-tagged canonical *Arp2* (*D. pseudoobscura*), Arp2D (*D. pseudoobscura*), or Arp2D2 (*D. auraria*) (see Figure 2A for design of transgenic lines). Native GFP fluorescence of the tagged proteins was imaged with actin and DNA probes. White boxes indicate the location of the enlarged image, and arrows indicate the cell cortex. Scale bars are 25 μm. **C)** RT-PCRs assessing expression of canonical *Arp2, Arp2D*, and codon-optimized *Arp2D* (see Figure 3A for design of transgenic lines). Females were dissected, separating carcass (‘C’) and ovaries (‘O’). *rp49* is shown to compare overall cDNA levels of the samples. The bottom gel image indicates cDNA samples generated without RT. No genomic DNA contamination was detected. RNA expression of *Arp2D* and codon optimized *Arp2D* were both detected at comparable levels. **D)** Schematic for a female fertility assay. Arp2- and codon optimized Arp2D-expressing females were crossed to Oregon R (wildtype) males. Total adult progeny is shown, indicating codon-optimized Arp2D still leads to female subfertility.

**Figure S4:**
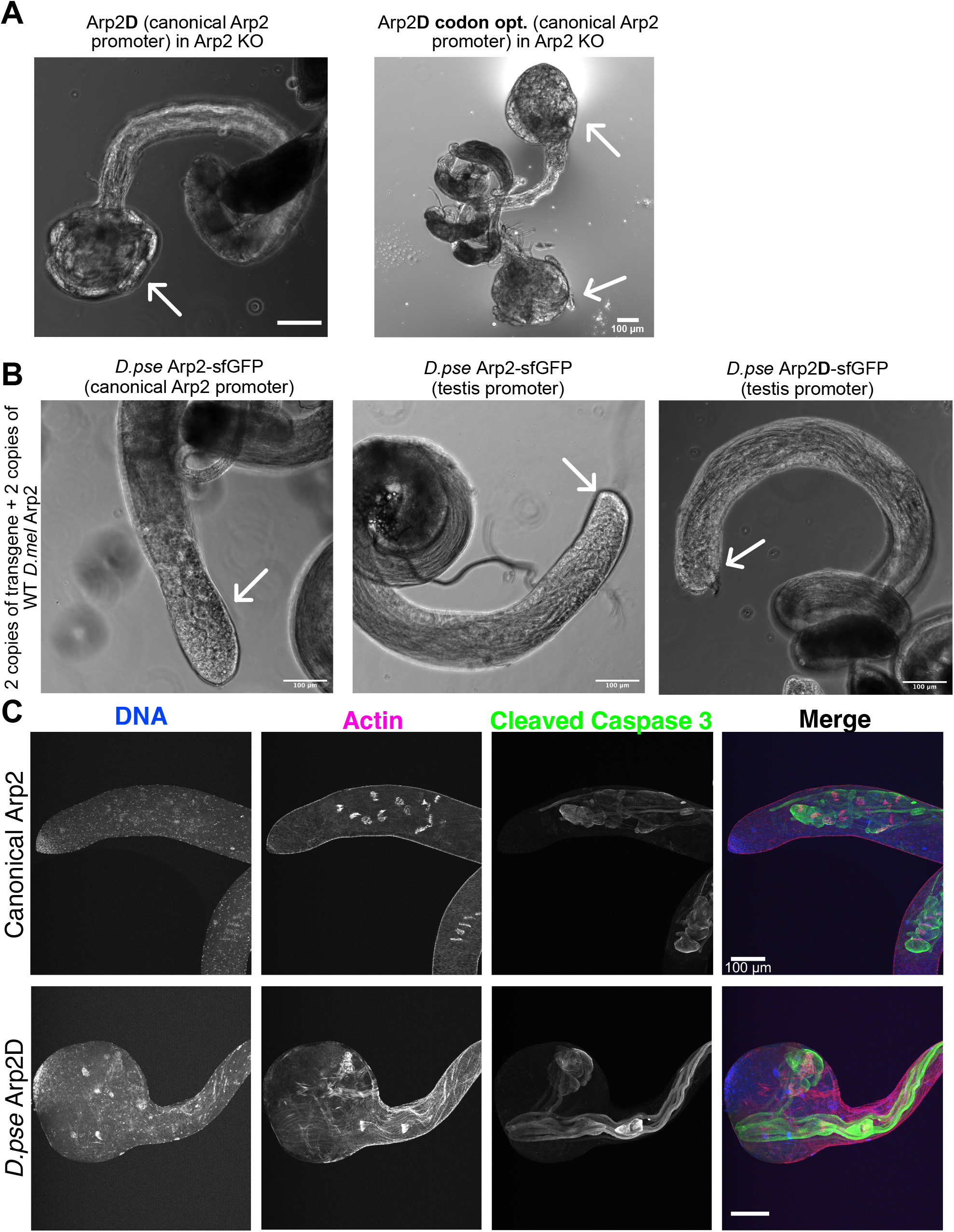
The testis morphology defect does not appear to be due to codon usage, overexpression, or lack of caspase activity. **A)** Brightfield images of testes from *Arp2*-KO flies expressing *D. pseudoobscura Arp2D* or codon-optimized *Arp2D*. Both genotypes exhibited enlarged testis apical ends, indicated with arrows. **B)** Brightfield images of testes from *D. melanogaster* flies that were modified to express sfGFP-tagged *D. pseudoobscura* Arp2 or Arp2D under the canonical Arp2 promoter or a promoter of a gene highly expressed in the testis (*Arp53D*). Despite two copies of the transgene and two copies of endogenous *Arp2*, testes appeared wildtype in morphology at the apical end (indicated by arrows), suggesting overexpression does not lead to a bulbous end. **C)** Immunofluorescent images of testes from *Arp2*-KO flies expressing canonical Arp2 or *D. pseudoobscura* Arp2D. DNA, actin, and cleaved caspase 3 (Asp 175), which indicates caspase activation, were imaged. As expected for normal sperm development, Arp2D-expressing testes exhibited caspase activity during individualization, similar to flies expressing canonical Arp2. All scale bars are 100 μm.

**Figure S5:**
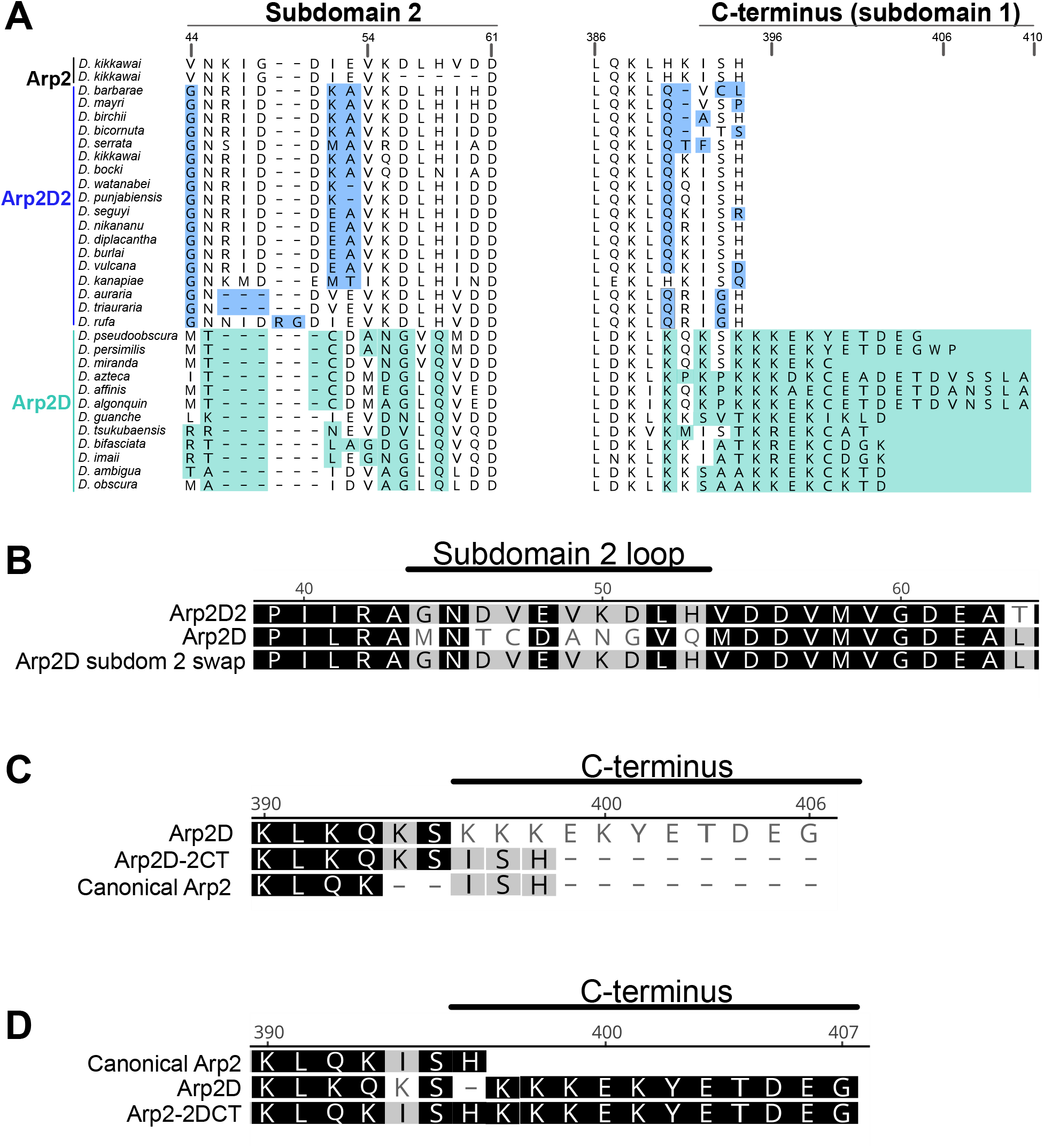
Sequence modification of Arp2D and canonical Arp2. **A)** Protein alignment of subdomain 2 and the C-terminus of Arp2, Arp2D2 and Arp2D. Both splice variants of Arp2 are shown. Significant regions of divergence are highlighted (teal for Arp2D and blue for Arp2D2). Numbers above the alignment are with respect to the canonical Arp2 protein sequence. **B)** A protein alignment of Arp2D2, Arp2D and the Arp2D chimera are shown. The chimera is Arp2D with *D. auraria*’s subdomain 2 (‘Arp2D subdom 2 swap’). **C)** The protein sequences of Arp2D, Arp2D-2CT, and canonical Arp2 are aligned. The C-terminus of Arp2D was replaced with Arp2’s short C-terminus, which is predicted to be disordered. **D)** Canonical Arp2, Arp2D, the chimera Arp2-2DCT are aligned, showing the C-terminus of Arp2D was added to canonical Arp2.

**Figure S6:**
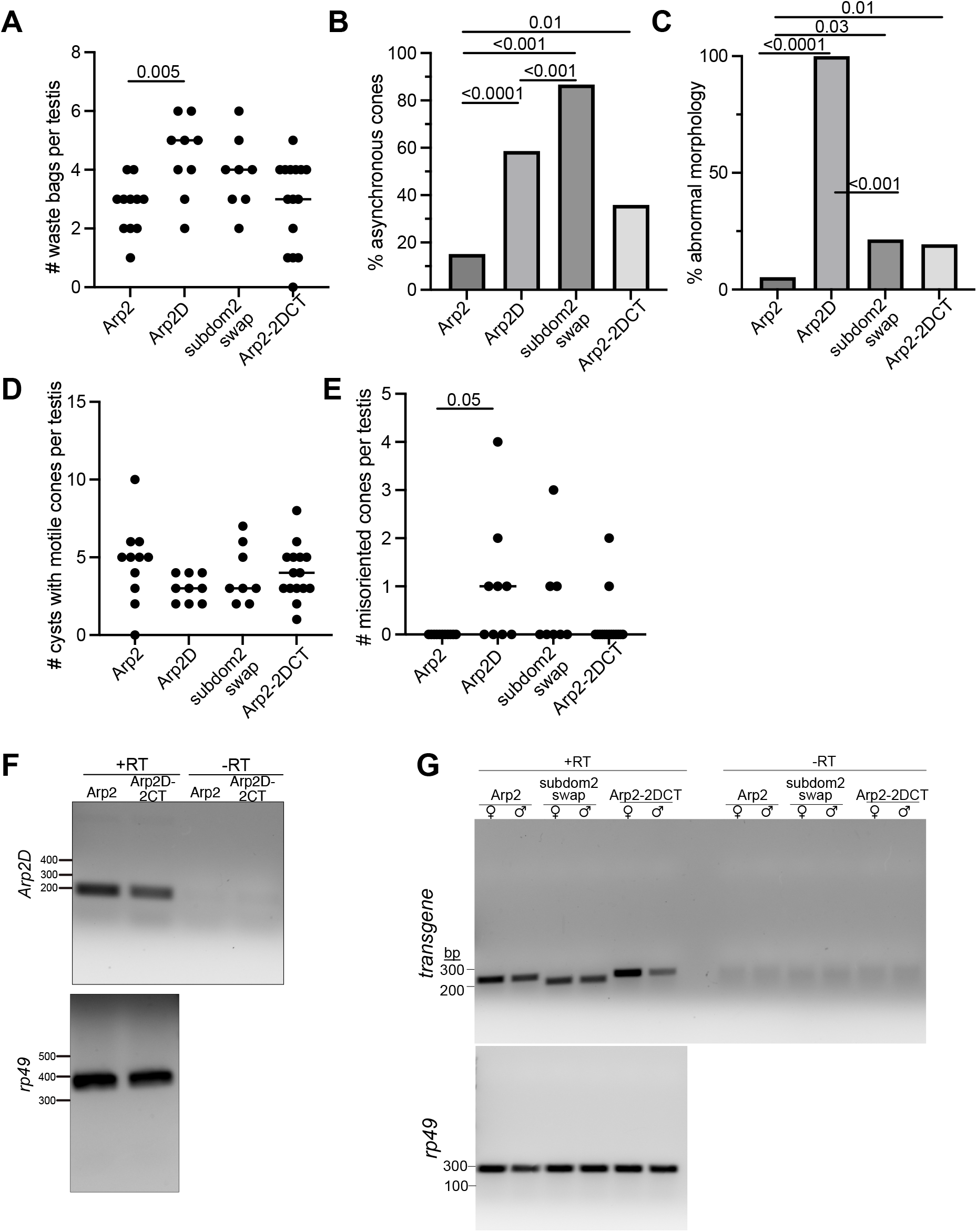
Transgene expression and quantification of testis cytology for Figures 4-6. **A-E)** Quantification of individualization defects. *Arp2*-KO flies expressing untagged canonical Arp2, Arp2D, Arp2D with modified subdomain 2 (‘subdomain2 swap’), and canonical Arp2 with Arp2D’s C-terminus (‘Arp2-2DCT’) were imaged and the following features were quantified: the number of waste bags per testis (A), the percent of cysts with asynchronous cones (B), the percent of testes with abnormal apical ends (C), the number of cysts with motile cones (no longer at sperm nuclei) per testis (D), and the number of misoriented cones (directed toward the basal end) per testis (E). Results with statistical significance are displayed in Figures 4-6. **F)** RT-PCRs assessing expression of *Arp2D* versus *Arp2D-2CT* in heterozygous whole female flies. Expression of *rp49* shows relative cDNA levels between samples, and ‘-RT’ samples (no reverse transcriptase) indicate absence of genomic DNA. **G)** RT-PCRs assessing expression of canonical Arp2, Arp2D subdomain 2 swap, and Arp2-2DCT in *Arp2*-KOs (left side of top image). Whole females (homozygous) and males (hemizygous) were analyzed separately. Genomic DNA was absent (‘-RT’, right side of top image) and cDNA levels were comparable among samples (*rp49* control, bottom image).

## Supplemental Information

**Data S1: Species and Primers**

Cultured species are listed, and scaffolds and accession numbers are provided for *Arp2* and *Arp2D2* sequences obtained from BioProject ID 554346^26^. All primers used for targeted sequencing of *Arp2D2* loci and RT-PCRs are also provided.

**Data S2: Sequences**

*Arp2D2* sequences obtained from *montium* species with unsequenced genomes are provided in addition to *Arp2* and *Arp2D2* coding sequences from BioProject ID 554346^26^ and annotated genomes^24,61^.

**Data S3: Tree**

The tree for Figure 1B is provided as a newick file with bootstrap support.

**Data S4: Constructs**

All construct sequences used for generating fly transgenics are provided.

